# *Mycobacterium abscessus* promotes *Pseudomonas aeruginosa* biofilm formation and antibiotic tolerance

**DOI:** 10.1101/2024.12.31.630905

**Authors:** Melissa S. McDaniel, Sara E. Edmonds, Evani N. Patel, Joshua J. Baty, Jessica A. Scoffield

## Abstract

Modulator therapies have improved outcomes for people with Cystic Fibrosis (pwCF), and currently more than 50% of pwCF are over the age of 18. This has resulted in an increased prevalence of atypical pathogens, including non-tuberculous mycobacteria (NTM). CF-isolation rates of NTM and *Pseudomonas aeruginosa* (*Pa*) are high, and those co-colonized have worse clinical outcomes. We therefore investigated the behavior of these two organisms in a dual-species biofilm. We found that coculture of *Mycobacterium abscessus* (MAB) promoted biofilm formation by *Pa*. Confocal imaging revealed changes in biomass and structural organization of the *Pa* biofilm during coculture with MAB. DNase treatment slightly decreased dual-species biofilm, but biofilm formation was completely abrogated in Pel- and Psl-deficient mutants of *Pa*. Moreover, dual-species cultures promoted tolerance of *Pa* to tobramycin treatment. Overall, our findings highlight an interaction between *P. aeruginosa* and *M. abscessus* that may result in bacterial persistence for pwCF during antibiotic therapy.

## INTRODUCTION

Cystic fibrosis (CF) is a genetic disorder caused by mutations in the cystic fibrosis transmembrane conductance regulator (CFTR). Pulmonary manifestations are characterized by hyperinflammation, muco-obstruction, and decreased mucus clearance, coupled with innate immune dysfunction^1–3^. These factors lead to chronic pulmonary bacterial infections that cannot be resolved, despite dramatic inflammation of the airways. Prior to the advent of CFTR modulator therapies, these infections and the associated decline in lung function were the primary drivers of morbidity and mortality in persons with CF (pwCF), and although these treatments have gone very far in improving lung function and quality of life, they have failed to completely eradicate bacterial infections in pwCF^4–8^. Further, the increasing age of pwCF and the long-term use of antibiotics to treat infections in this population has led to the emergence of non-traditional and antibiotic-resistant CF pathogens, including non-tuberculous mycobacteria (NTM)^9^.

NTM are a diverse group of bacteria commonly isolated from environmental sources that can act as opportunistic pathogens, causing a variety of disease manifestations within chronic respiratory infections^10,11^. In pwCF, *Mycobacterium abscessus* (MAB) and *Mycobacterium avium* complexes (MAC) are the most prevalent NTM species, with MAB colonization being associated with a more severe decline in lung function as compared to MAC or other traditional CF pathogens^12,13^. Unlike infections with *Mycobacterium tuberculosis,* NTM infections may exist both intracellularly (with macrophages as the primary reservoir), or extracellularly, likely as a biofilm within the lung^14,15^. This makes the ability to persist within a biofilm potentially important for the virulence of this organism, both for the mechanism of initial acquisition and for persistence in the lung^14–18^. MAB also has two known morphotypes, with the smooth (S) morphotype indicating production of glycopeptidolipids, and the rough (R) morphotype associated with the loss of their production^19,20^. The rough morphotype is known to emerge over the course of human disease, increases resistance to phagocytosis, and is associated with a more virulent infection^19,21–27^.

Thickened mucus and a decrease in mucociliary clearance predisposes the CF lung to colonization with pathogens in the form of a chronic, complex polymicrobial biofilm, in which the predominant pathogen is *Pseudomonas aeruginosa* (*Pa*). This highly virulent organism is found in over half of pwCF^28^, and is known to interact with many CF pathogens during co-infection to alter infection dynamics^29–44^. *P. aeruginosa* also displays well-studied adaptations specific to the CF lung, including the transition from early exopolysaccharides such as Pel and Psl, to the late-stage exopolysaccharide alginate. Each of these exopolysaccharides have been shown to shape interactions with other organisms, particularly with the common CF pathogen *Staphylococcus aureus*^42,45–51^.The acquisition of NTM is not well correlated with the presence of *Pa,* although they can be co-isolated at relatively high rates^52–54^. Importantly, non-CF bronchiectasis patients co-infected with NTM and *Pa* are well-documented to have a steeper rate of lung decline than those patients infected with either alone, indicating synergy between these two organisms in terms of virulence^55–57^. In further support of this, previous publications that focused on dual-species interactions between *Pa* and MAB have indicated that co-culture can provide a benefit to both species. MAB and *Pa* can form dual-species biofilms, and co-culture in the presence of antibiotic treatment promotes MAB growth^58,59^. It has also been shown that MAB can directly degrade the PQS quorum-signaling molecule of *Pa* via a dioxygenase (Aqd). However, these studies did not directly the address the role of MAB on *Pa* physiology and fitness^60–62^.

Previous experimental and epidemiological studies indicate that interactions between *Pa* and NTM contribute to severity of infection. We therefore decided to investigate the behavior of these two organisms during co-culture in a dual-species biofilm. We found that despite rapid killing of MAB during co-culture, overall biofilm formation by *P. aeruginosa* increased in a manner dependent on both extracellular DNA (eDNA) and the Psl and Pel polysaccharides of *P. aeruginosa*. Co-culture of these two organisms also increased tolerance of *P. aeruginosa* to tobramycin, an anti-Pseudomonal antibiotic commonly used in a CF context. Ultimately, these results indicate that dual-species interactions between *P. aeruginosa* and MAB may contribute to an increase in disease severity during pulmonary infection.

## RESULTS

### Coculture of *P. aeruginosa* and MAB promotes biofilm formation

To investigate the impact of dual-species co-culture on biofilm formation by *P. aeruginosa* and MAB, we first grew single- and dual-species biofilms with *P. aeruginosa* PAO1, a standard non-mucoid lab strain or *P. aeruginosa* mPA08-31, a mucoid clinical strain (Supplementary Figure 1A), with *M. abscessus* smooth or rough morphotypes (Supplementary Figure 1B) at varying time intervals. Biofilm biomass was measured via crystal violet assay normalized to total growth. After 24 hours, biofilm biomass significantly increased when *P. aeruginosa* PAO1 was co-cultured with either the smooth or rough morphotypes of MAB (*p* < 0.0001, *p* = 0.0125). However, this phenotype was lost at the 48- and 72-hour time points (Figure 1A). Conversely, when *P. aeruginosa* mPA08-31 was co-cultured with smooth or rough MAB, there was no difference between single- and dual-species biofilms at 24 hours post-inoculation. However, an increase in overall biomass was observed with the smooth and rough morphotype at 48 hours post-inoculation (*p* = 0.0032, *p* < 0.0001), and with only the rough morphotype at 72 hours post inoculation (*p* = 0.0012) (Figure 1B). To determine which species was responsible for the increase in biofilm, we then measured viable colony forming units (CFUs) of each species via a differential plating scheme. There was no significant increase in *P. aeruginosa* CFUs for either strain compared to single species culture at any of the time points measured except for the 72-hour time point for mPA08-31 with the rough morphotype (*p =* 0.004) (Figure 1C,D). As expected from known patterns of aggregation in MAB, there were significantly fewer viable CFUs of the rough morphotype of MAB than of the smooth in a single species biofilm at all time points tested (Figure 1E, F). During co-culture with either *P. aeruginosa* PAO1 or mPA08-31, viable MAB rapidly decreased, with an approximately 3-log drop in CFUs of MAB(S) by 48 hours post-infection. With PAO1, MAB was below 10^2^ CFU/mL (the limit of detection) by 72-hours (Figure 1E), while ∼10^4^ CFU/mL of MAB remained when co-cultured with mPA08-31 (Figure 1F), possibly explaining differences in kinetics of the cooperativity. Taken together, these data indicate that for several strains of *P. aeruginosa,* co-culture with MAB results in an increase in total biofilm biomass, despite a dramatic decline in viable MAB.

**Figure 1.**
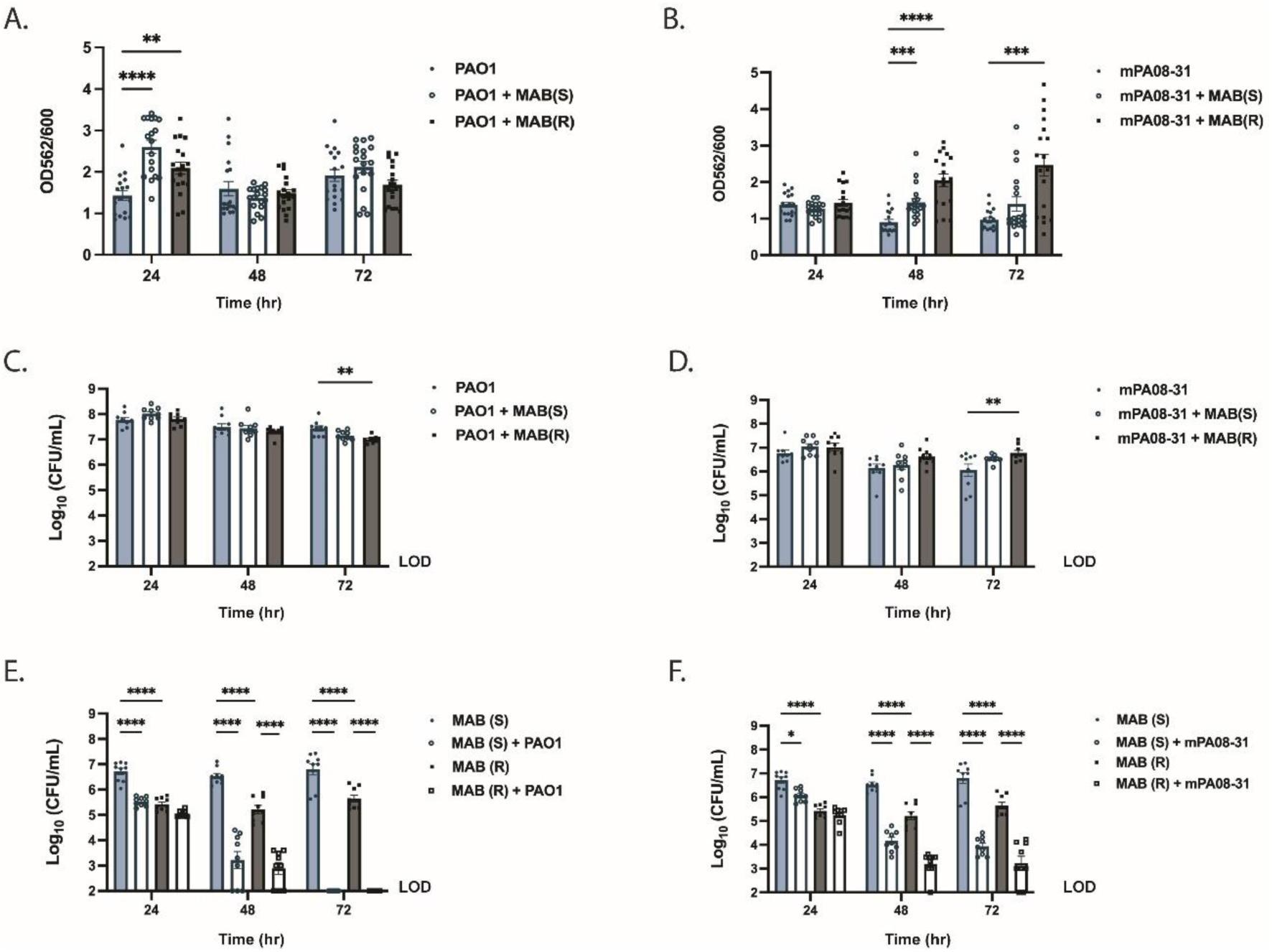
Co-culture of *P. aeruginosa* and MAB promotes biofilm formation. *M. abscessus* ATCC 19977 smooth and rough morphotypes were co-cultured with either of two *P. aeruginosa* strains (PAO1 or mPA08-31) in LB medium in a 96-well plate for 24, 48, or 72 hours at 37 °C. Biofilm biomass was then measured using crystal violet staining for A) PAO1 or B) mPA08-31 during single- and co-culture (n = 3 biological replicates, 6 technical). Two-way ANOVA with Tukey’s multiple comparisons test. Viable colony forming units of *P. aeruginosa* C) PAO1 or D) mPA08-31 during single- and co-culture, and of *M. abscessus* during single- and co-culture with E) PAO1 and F) mPA08-31 were measured (n = 3 biological replicates, 3 technical). Two-way ANOVA with Tukey’s multiple comparisons test.

Having observed an increase in biomass via crystal violet assay, we then investigated the biofilm structure using confocal microscopy. Single- and dual-species biofilms with *P. aeruginosa* PAO1^GFP+^ and MAB^mCherry+^ smooth or rough morphotypes were grown for 24, 48, or 72 hours, and the biofilm structure was subsequently imaged via confocal microscopy. As expected from the crystal violet assay, dual-species biofilms were visibly thicker than single-species biofilms with *P. aeruginosa* PAO1 alone at the 24-hour time point (Figure 2A). Interestingly, we observed that MAB was localized primarily to the bottom of the biofilm, whereas *P. aeruginosa* can be found at the top, a phenomenon that has been seen by other groups^59^. To confirm our visual findings, we quantified both biofilm height and volume for each species separately using BiofilmQ. We found that PAO1 biofilm volume was significantly increased during co-culture with both morphotypes of MAB at 24 hours (*p* = 0.0182, *p* = 0.0079), but not at subsequent time points (Figure 2B, Supplementary Figure 2A). PAO1 biofilm height was only significantly increased with MAB(S) at the 24-hour time point (p = 0.0455), and only with MAB(R) at the 72-hour time point (p = 0.0044) (Figure 2C, Supplementary Figure 2C). MAB biofilm volume trended towards a decrease in biomass as compared to single species biofilms at 24 hours, but was significantly decreased at 48 and 72 hours (Figure 2D, Supplementary Figure 2B). No significant changes in biofilm height were seen for MAB (Figure 2E, Supplementary Figure 2D). To quantify differences in localization within the dual species biofilm, we plotted distribution of distance to substrate for each cell within the biofilm. At 24 hours, the distribution of *P. aeruginosa* was notably right shifted compared to the smooth morphotype of *M. abscessus* (Figure 2F). However, species were more evenly distributed when P. aeruginosa was grown with the rough morphotype of *M. abscessus* (Figure 2G) with similar trends at all timepoints tested (Supplementary Figure 2E,F). We repeated these experiments for *P. aeruginosa* mPA08-31. Like with PAO1, we saw a visible increase in mPA08-31 biofilm thickness during co-culture, although this was not statistically significant when quantified apart from biofilm height when co-cultured with the rough morphotype at 72 hours (*p =* 0.0079) (Supplementary Figure 3A-C). In these biofilms MAB volume decreased significantly during co-culture only with the smooth morphotype (Supplementary Figure 3D, E). The relative distance to substrate was also like that seen in PAO1, though differences in localization were strongest at the 48- and 72-hour time points with the smooth morphotype (Supplementary Figure F, G).

**Figure 2.**
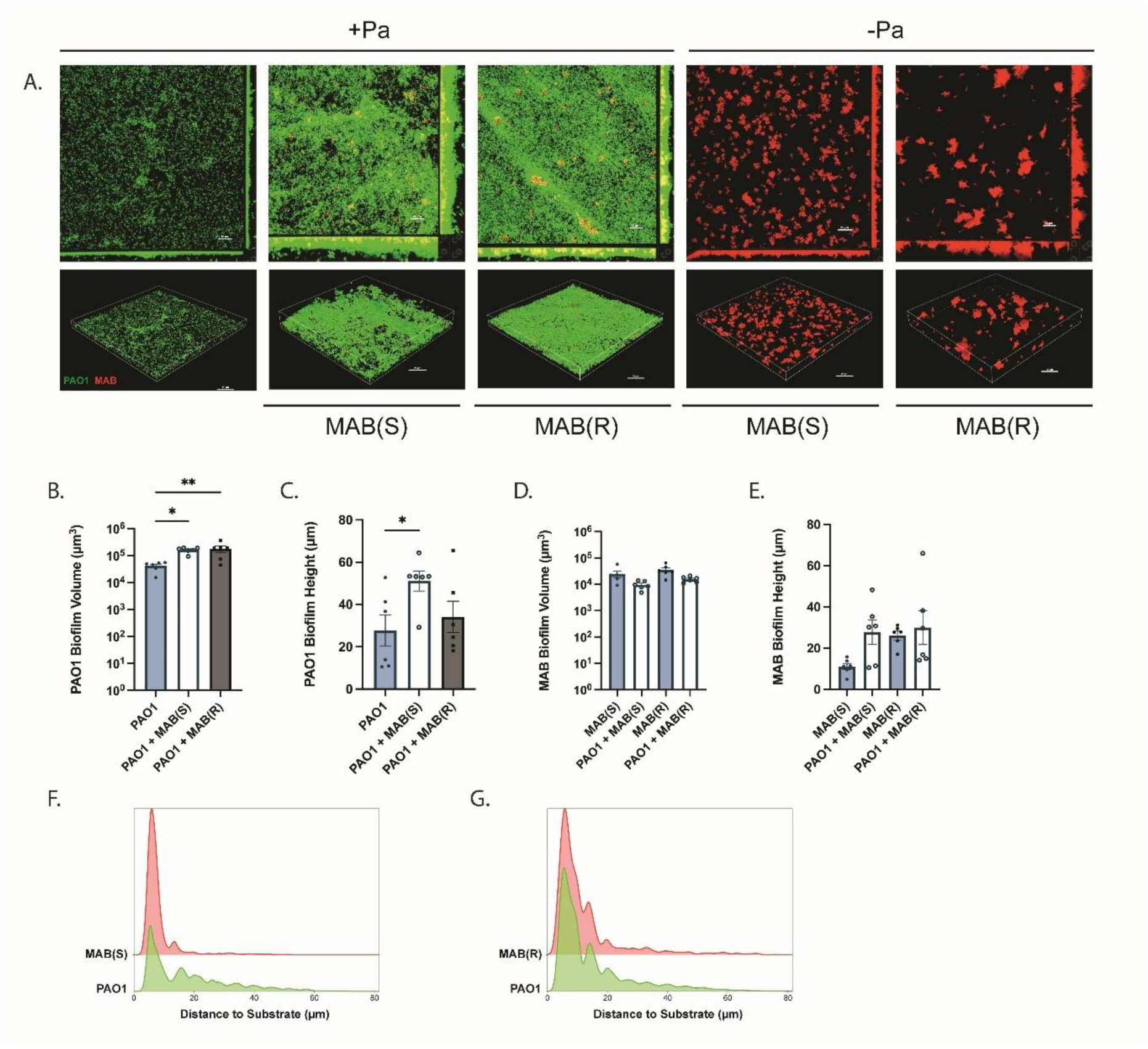
Imaging of dual species *P. aeruginosa* and MAB biofilms shows changes in topology during co-culture. Fluorescently labeled *M. abscessus* ATCC19977 smooth and rough morphotypes (mCherry+) were co-cultured with *P. aeruginosa* PAO1 (GFP+) in LB medium in an 8-well μSlide for 24 hours at 37 °C (n = 2 biological replicates, 3 technical). A) Structural composition of single- and dual-species biofilms was evaluated via confocal imaging at 40X magnification. B) Volume and C) height of *P. aeruginosa* PAO1 and D) volume and E) height of *M. abscessus* in single- and dual-species biofilms were quantified via BiofilmQ^81^. One-way ANOVA with Tukey’s multiple comparisons test. Histogram of pixel distribution of *P. aeruginosa* PAO1 and *M. abscessus* distance to substrate for F) smooth and G) rough morphotypes in dual-species biofilms. (n = 2 biological replicates, 3 technical).

### Extracellular DNA is not the primary contributor to increased biofilm formation

Due to the increase in overall biomass in our dual-species biofilms without a large increase in viable CFUs of *P. aeruginosa,* but with a dramatic decrease in live MAB, we hypothesized than an extracellular matrix component was likely responsible for the increase in crystal violet staining. We also noted that similarities in kinetics of MAB killing by *P. aeruginosa* and the times at which biofilm were increased. We hypothesized that extracellular DNA (eDNA) from dead MAB might be responsible for the increased biomass in the dual-species biofilm. To test this, we grew single- or dual-species biofilms with or without DNase treatment and measured total biomass via crystal violet staining. For *P. aeruginosa* PAO1, crystal violet staining of dual-species biofilms without DNase treatment produced similar results to those in Figure 1, with a significant increase in total biomass seen with dual-species biofilms only at the 24-hour time point and primarily for MAB(S) (p *=* 0.0069). With DNase treatment, we observed reduced biomass at all timepoints, indicating the presence of eDNA in the biofilm. However, this did not seem to impact the observed dual species phenotype at the 24h timepoint, and appeared to increase the dual species phenotype at the 48- and 72-hour time points for both morphotypes. These timepoints showed significantly increased biofilm during co-culture for MAB(R) (*p* = 0.0319, *p* = 0.0469) (Figure 3A), indicating that the dual species biofilm is less dependent on eDNA at the later time points. For mPA08-31, we again see a significant increase in biofilm biomass for both MAB morphotypes at 48- (*p* = 0.0467, *p* = 0.0049) and 72-hours (*p =* 0.0332, *p =* 0.0470). With DNase treatment, this increase was completely gone at 48-hours, but biomass was significantly increased for MAB(R) co-culture at 72 hours (*p* = 0.0026) (Figure 3B). These results may indicate a *P. aeruginosa* strain-specific biofilm composition change in dual-species cultures.

**Figure 3.**
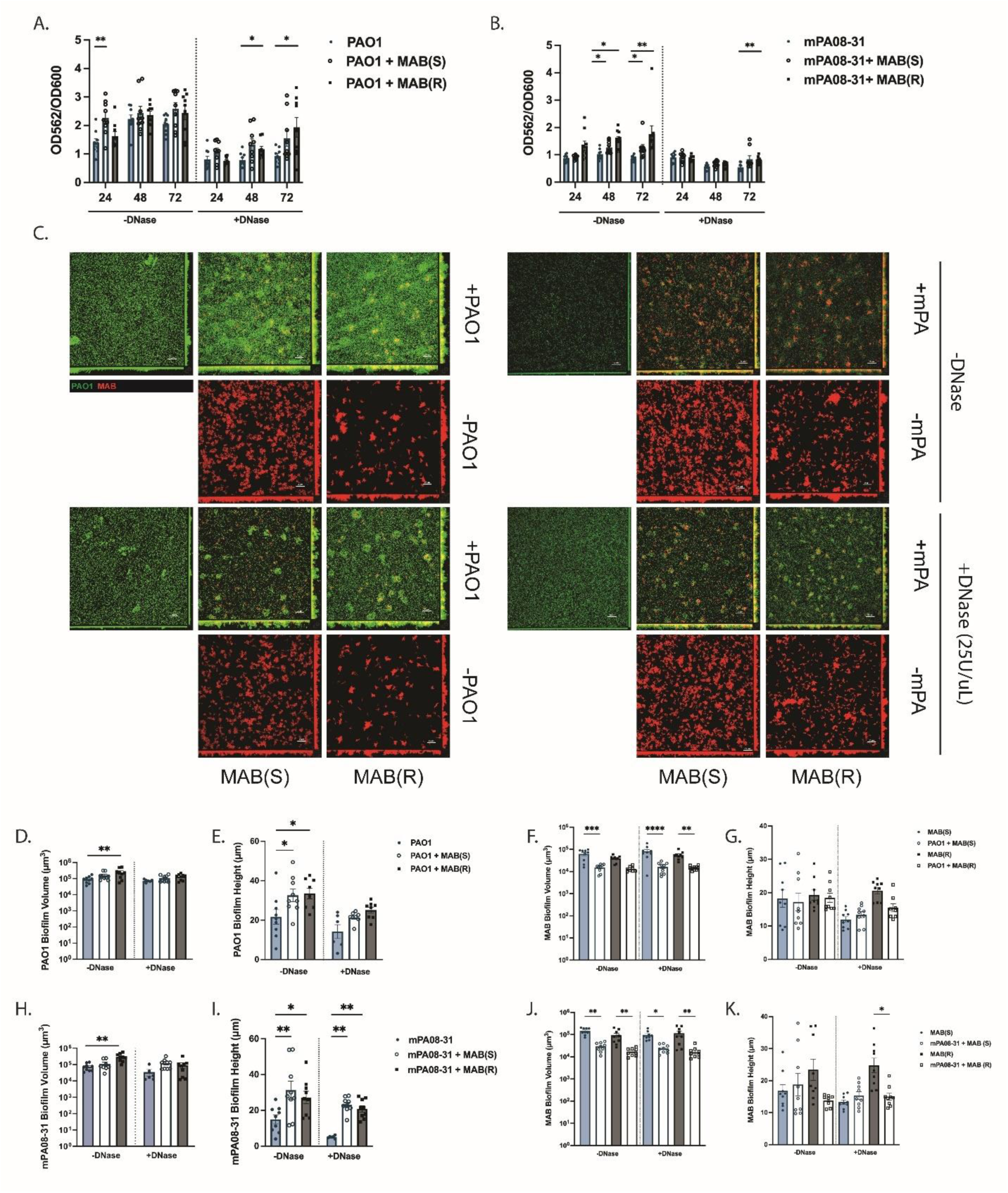
DNase treatment partially abrogates dual-species cooperativity between *P. aeruginosa* and MAB. *M. abscessus* ATCC 19977 smooth and rough morphotypes were co-cultured with either of two *P. aeruginosa* strains (PAO1 or mPA08-31) in LB medium in a 96- well plate for 24, 48, or 72 hours at 37 °C. Biofilms were then treated for 1 hour with DNase (25U/mL) or PBS. Biofilm biomass was then measured using crystal violet staining for A) PAO1 or B) mPA08-31 during single- and co-culture (n = 3 biological replicates, 3 technical). Two-way ANOVA with Tukey’s multiple comparisons test. Fluorescently labeled *M. abscessus* ATCC19977 smooth and rough morphotypes (mCherry+) were co-cultured with *P. aeruginosa* PAO1^GFP+^ in LB medium for 24 hours or with *P. aeruginosa* mPA08-31^GFP+^ for 72 hours in an 8-well μSlide at 37 °C (n = 2 biological replicates, 3 technical). C) Structural composition of single- and dual-species biofilms with and without DNase treatment was evaluated via confocal imaging at 40X magnification. D,E) Volume and height of *P. aeruginosa* PAO1 and F,G) *M. abscessus* or H,I) *P. aeruginosa* mPA08-31 and J,K) *M. abscessus* single- and dual-species biofilms were quantified via BiofilmQ^81^ (n = 3 biological replicates, 3 technical). Kruskal-Wallis with Dunn’s multiple comparisons test.

To confirm the crystal violet results, we performed confocal microscopy on single- and dual-species biofilms of *P. aeruginosa* PAO1 after 24 hours, and *P. aeruginosa* mPA08-31 after 72 hours, with or without a subsequent 1-hour DNase (25U/mL) treatment. Without DNase treatment, PAO1 biofilms were thicker and denser with either MAB(S) or MAB(R) present. With DNase treatment, dual-species biofilms, although still visibly taller, appeared to have reduced density compared to those without DNase treatment. Similar trends were also seen for biofilms grown with mPA08-31, though there was perhaps less of an effect of DNase on biofilm composition for this strain (Figure 3C). These results were confirmed through quantification, where PAO1 biofilm volume trended higher with MAB(S) and was significantly increased with MAB(R) without DNase treatment (*p* = 0.0030), and the significance of this increase was lost with DNase treatment (Figure 3D). PAO1 biofilm height was significantly increased with either MAB(S) or MAB(R) without DNase treatment (*p* = 0.0233, *p =* 0.0154). This significance was lost following DNase treatment, although both groups still trended higher than the single species biofilm (Figure 3E). For mPA08-31, biofilm volume trended higher with MAB(S) and was significantly increased with MAB(R) without DNase treatment (*p* = 0.0058), with the significance again lost with DNase treatment (Figure 3H). mPA08-31 biofilm height was significantly increased with either MAB(S) or MAB(R) without DNase treatment (*p* = 0.0012, *p* = 0.0204) but remained significant after DNase treatment (*p =* 0.0014, *p =* 0.0061) (Figure 3I). Taken together, these data indicate that eDNA is an important component of both single and dual-species biofilms but is likely not the major driver of the increase in biofilm formation.

### Both Psl and Pel contribute to increased biofilm biomass

To investigate what other factors might be responsible for the increase in biofilm biomass during dual-species co-culture, we examined the contribution of major exopolysaccharides of *P. aeruginosa.* Because we saw an increase in biofilm formation for both mucoid and non-mucoid strains of *P. aeruginosa*, we thought it was unlikely that alginate was responsible for our dual-species phenotype. We therefore tested PAO1 mutants deficient in Pel and Psl, which are exopolysaccharides that are predominantly produced by non-mucoid isolates. Moreover, it is known that Pel crosslinks with eDNA, which could potentially explain the loss of dual biofilm formation in DNase treated dual biofilms with PAO1. We grew single- and dual-species biofilms with parent PAO1 or with transposon mutants of PAO1 for *pslA*, *pelA*, or both *pslA* and *pelA* and evaluated our established cooperative phenotype via crystal violet assay. With the parent PAO1 we observed the expected significant increase in biomass at 24 hours with MAB(S) or MAB(R) (*p* = 0.0008, *p* = 0.0076) (Figure 4A). With the disruption of Pel this increase was lost at 24 hours, but a significant increase in biofilm biomass with the addition of MAB(S) and MAB(R) emerged at the 72-hour time point (*p =* 0.0390, *p =* 0.0064) (Figure 4B). With the disruption of Psl we observed a substantial reduction in overall biomass as expected, however dual species biofilm biomass trended higher with the MAB(S) morphotype and was significantly higher with MAB(R) at 24-hours (*p* = 0.0355). This increase was absent at 48- and 72-hour time points, like in the parent strain (Figure 4C). With disruption of both Psl and Pel, there was very little biomass detected and we saw complete abrogation of the phenotype, with no significant increase in biomass with either morphotype of MAB at any of the time points tested (Figure 4D). To rule out the possibility that changes to dual-species cooperativity were due to changes in the ability of mutant strains to kill MAB, we measured viable colonies for each at 24, 48, and 72-hours. No defect in killing was seen for any of the mutants tested with the exception of the *pslA pelA* double mutant at the 48-hour time point when cocultured with MAB(S) (*p =* 0.0367) (Supplementary Figure 6A,B). Mucoid strains of *P. aeruginosa,* such as mPA08-31, primarily produce alginate and may show variable amounts of Pel production^63,64^. We therefore measured Pel production via Congo Red assay and confirmed that *P. aeruginosa* mPA08-31 is capable of producing Pel, particularly when grown at 37°C (Supplementary Figure 5). Lastly, we grew dual-species biofilms with MAB and either *P. aeruginosa* PA14 or FRD1. PA14 produces Pel but not Psl or alginate, and we found an increase in biofilm biomass during co-culture with MAB, again indicating that either Pel or Psl, but not both, are required for the increased biomass during co-culture with MAB (Supplementary Figure 4B,D,F). FRD1, however, produces mainly alginate, and we observed only a slight increase in biomass at 24 hours co-culture with MAB(R) (Supplementary Figure 4A,C,E). Taken together, these data indicate a particularly important role for Pel in early biofilm increase during co-culture with MAB, though a compensatory mechanism involving Psl may exist at later timepoints.

**Figure 4.**
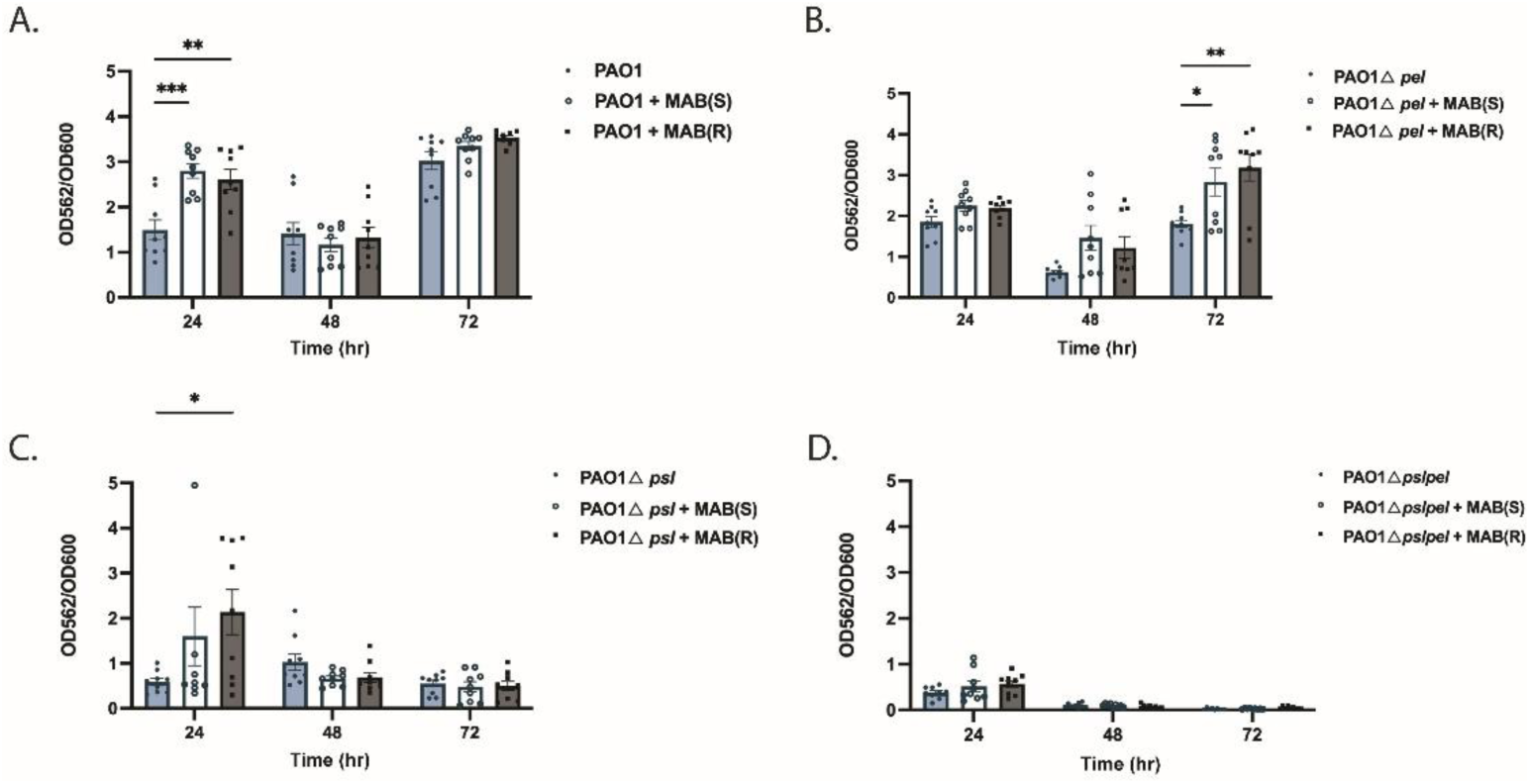
The Pel exopolysaccharide contributes to the increase in biofilm biomass during co-culture. *M. abscessus* ATCC 19977 smooth and rough morphotypes were co-cultured with *P. aeruginosa* strains PAO1, PAO1Δ*psl*, PAO1ΔΔ*pslpel* in LB medium in a 96-well plate for 24, 48, or 72 hours at 37 °C. Biofilm biomass was then measured using crystal violet staining for A) PAO1, B) PAO1Δ*pelA,* C) PAO1Δ*psl* or D) PAO1ΔΔ*pslpel,* during single- and co-culture (n = 3 biological replicates, 3 technical). Two-way ANOVA with Tukey’s multiple comparisons test.

### Dual-species biofilms promote tolerance of *P. aeruginosa* to tobramycin

The Pel and Psl exopolysaccharides are both known to contribute to resistance to antimicrobials such as tobramycin, which is often used in CF-related *P. aeruginosa* infections^65–68^. We therefore decided to test the susceptibility of *P. aeruginosa* to tobramycin treatment in a single-species biofilm as compared to in a dual-species biofilm with MAB. We grew single- and dual-species biofilms as in previous experiments, and then replaced media with either PBS or tobramycin (50ug/mL) for 1 hour. As expected from previous experiments, we saw no difference in *P. aeruginosa* CFUs with or without MAB present. With tobramycin treatment there was a decrease in live *P. aeruginosa* when by itself, but a lesser decrease in live *P. aeruginosa* with MAB present, with a significant difference between tobramycin treated single-species culture and dual-species culture with MAB(S) for PAO1 (*p =* 0.0021), and both morphotypes for mPA08-31 (*p =* <0.0001, *p* = 0.0003) (Figure 5A, B). Interestingly, this tolerance for mPA08-31 did not extend to the 48- and 72-hour time points for which we had previously seen the biggest difference in biofilm formation, perhaps indicating that a second mechanism is at play (Supplementary Figure 7B). Changes in *P. aeruginosa* burden did not lead to any subsequent differences in the amount of *M. abscessus* present in dual-species biofilms with PAO1 or mPA08-31 (Supplementary Figure 7A,C). To observe the effect of coculture on susceptibility of *P. aeruginosa* to tobramycin in terms of biofilm structure, we again performed confocal microscopy on single- and dual-species biofilms of *P. aeruginosa* PAO1 after with or without a subsequent 1-hour tobramycin treatment (50ug/mL) treatment. As expected from the CFU results, dual-species biofilms show significantly increased *P. aeruginosa* volume (*p =* 0.0096) (Figure 5C, D), which became more dramatic with tobramycin treatment (*p =* 0.0054*, p =* 0.0041). These data indicate that in addition to thickening of the biofilm matrix, co-culture with MAB also decreases susceptibility of *P. aeruginosa* to the common anti-pseudomonal drug tobramycin.

**Figure 5.**
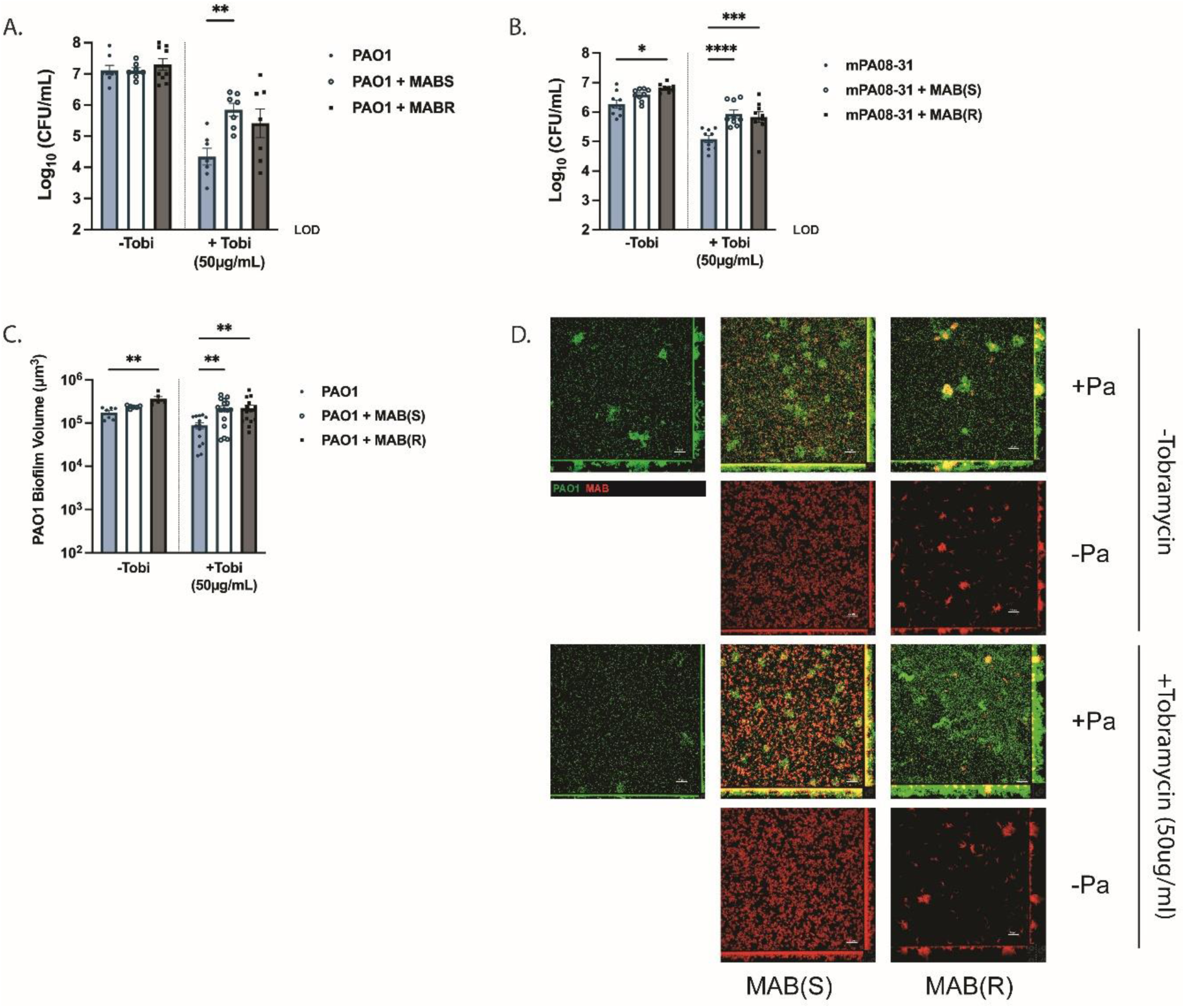
Co-culture of *P. aeruginosa* and MAB decreases susceptibility of *P. aeruginosa* to tobramycin. *M. abscessus* ATCC 19977 smooth and rough morphotypes were co-cultured with PAO1 or mPA08-31 in LB medium in a 96-well plate for 24 hours at 37 °C. Biofilms were then treated for 1 hour with tobramycin (50 μg/mL) or 1X PBS. Viable colony forming units of *P. aeruginosa* A) PAO1 and B) mPA08-31 during single- and dual-species culture with and without tobramycin treatment were measured via differential plating. (n = 3 biological replicates, 3 technical). One-way ANOVA with Tukey’s multiple comparisons test. Fluorescently labeled *M. abscessus* ATCC19977 smooth and rough morphotypes (mCherry+) were co-cultured with *P. aeruginosa* PAO1 (GFP+) in LB medium for 24 hours in an 8-well μSlide at 37 °C (n = 2 biological replicates, 3 technical). C) Volume and biofilms were quantified via BiofilmQ. Kruskal-Wallis with Dunn’s multiple comparisons test. D) Structural composition of single- and dual-species biofilms with and without tobramycin treatment was evaluated via confocal imaging at 40X magnification.

### Dual species coculture results in changes to expression of an aliphatic amidase

Lastly, we were interested in investigating the mechanism by which MAB induces an increase in biofilm formation by *P. aeruginosa.* We therefore performed RNA-sequencing on *P. aeruginosa* biofilms grown either alone or in conjunction with MAB. To harvest enough biofilm for adequate RNA quantities, biofilms were switched to a 6-well format, with each sample pooled between 3 wells at 48-hours (Supplementary Figure 8). RNA harvested from pooled biofilm samples was prepared for sequencing and sequenced at a read depth of ∼20 million reads per sample. Principal-component analysis of gene expression data indicated some differences in the presence of MAB (Figure 6A). Overall, differential expression analysis between samples showed that 57 genes are differentially regulated between dual species biofilms as compared to *P. aeruginosa* alone, with the top 20 represented in a heatmap (Figure 6B). The most highly differentially expressed genes included a downregulation of every gene in the aliphatic amidase operon, several genes of which are known to influence biofilm formation and other virulence phenotypes in *P. aeruginosa* (Figure 6C). To confirm these results, we measured expression of these genes in single and dual-species biofilms via qPCR, and again found a downregulation in the presence of MAB (Figure 6D). These results indicate that although coculture of *P. aeruginosa* with MAB does not result in global transcriptomic changes, differences in the expression of the aliphatic amidase operon are likely contributing to our cooperative dual-species phenotype. Further investigations into the role of this pathway during dual-species infections with MAB may help the development of antimicrobial strategies to polymicrobial infections, and aid in our understanding of the increase in infection severity seen in patients when both organisms are present.

**Figure 6.**
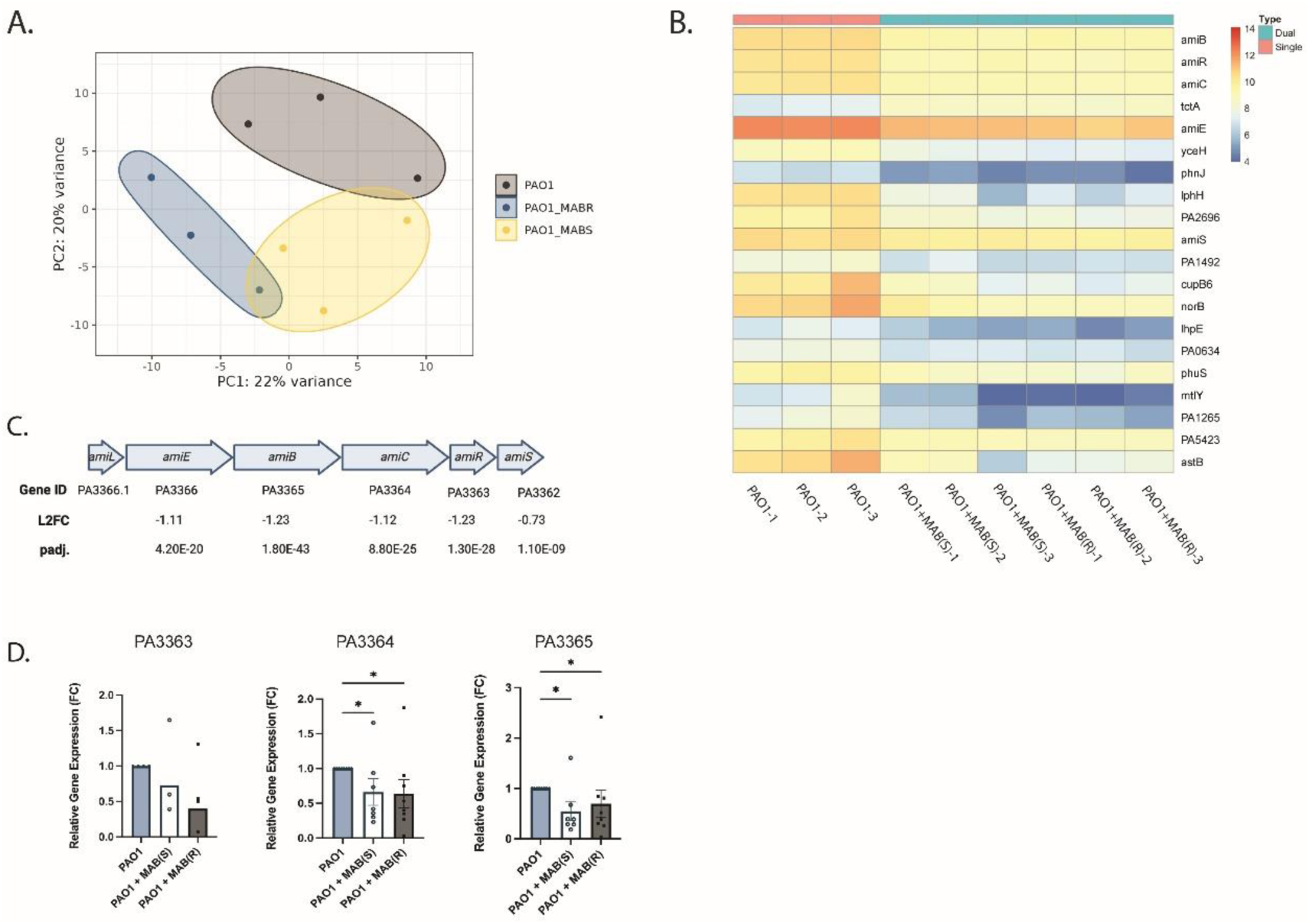
Co-culture of *P. aeruginosa* and MAB changes the transcriptomic profile of *P. aeruginosa.* *M. abscessus* ATCC 19977 smooth and rough morphotypes were co-cultured with *P. aeruginosa* PAO1 in LB medium in a 6-well plate for 48 hours at 37 °C. Adherent cells were pooled (3 per sample) for RNA extraction and sequencing (n = 3). A) Principal-component analysis of samples based on RNA sequencing data, colored by infection group. B) Heatmap of differentially expressed genes during *Pa* single or dual-species biofilms determined via differential expression analysis using DESeq2. C) Graphical representation of the aliphatic amidase operon of PAO1 with corresponding log2foldchange and padj. values for dual-species biofilms with either morphotype of MAB vs. single-species biofilms. D) Relative gene expression by PAO1 during single- or dual-species culture of genes identified via RNA-seq analysis as measured by RT-qPCR. Kruskal-Wallis with Dunn’s multiple comparisons test.

## DISCUSSION

In this study, we sought to explore interactions between *Pa* and MAB to better understand how they might contribute to severity of CF infection. We established a dual-species biofilm model where we measured biofilm biomass, architecture, and tolerance to antimicrobial treatment. Taken together, our results indicate that despite rapid killing of MAB during co-culture, overall biofilm biomass increased for a variety of *P. aeruginosa* strains used, and independent of MAB morphotype. This increase in biomass was not the result of a corresponding increase in bacterial cell number, therefore we investigated the contribution of biofilm matrix components. We found that DNase treatment partially abrogated the dual species increase in biomass, indicating that eDNA is likely involved, although this varied temporally and by *P. aeruginosa* strain. Disruption of the two major exopolysaccharides of non-mucoid *P. aeruginosa* strains, Psl and Pel, indicated that both polysaccharides are likely involved in the observed phenotype but may be involved at different timepoints, with Pel showing more of an effect at 24h while Psl appears more involved in later timepoints. The DNase treatment results mirrored those of the Pel disruption, suggesting that eDNA and Pel are collectively acting to increase dual-species biofilm biomass at early timepoints. Co-culture of these two organisms also increased tolerance of *P. aeruginosa* to tobramycin, an anti-Pseudomonal antibiotic commonly used in a CF context. Lastly, we performed transcriptome analysis and found that in the presence of MAB, *P. aeruginosa* differentially expresses an operon involved in production of aliphatic amidases which is known to influence a variety of virulence phenotypes. Ultimately, these results indicate that dual-species interactions between *P. aeruginosa* and MAB may contribute to an increase in disease severity during pulmonary infection, and to recalcitrance of infections even with appropriate antibiotic treatment.

Initially we found that co-culture of a variety of strains of *P. aeruginosa* with either the rough or smooth morphotypes of MAB resulted in a thicker biofilm than either of these species alone. Consistent with previous reports of *P. aeruginosa* and MAB co-culture, imaging of biofilm structure showed MAB primarily on the substratum, with *P. aeruginosa* localized throughout the biofilm^59^. This might indicate that MAB is protected by *P. aeruginosa* matrix from either antimicrobial treatment, oxidative stress, or phagocytosis, although we did not measure these directly. Cooperative dual-species interactions with an increase in total biomass are often the result of a promotion of growth for one or both species involved. However, we found that there was no significant increase in the growth of *P. aeruginosa* and a dramatic decrease in live MAB present during co-culture. This indicated that it was likely an extracellular matrix component that was responsible for the increase in biofilm formation. Because of the relative proportion of MAB as compared to *P. aeruginosa* in the biofilms at later time points and the increase in biomass regardless of the MAB morphotype, we felt that the matrix component was likely either being produced by *P. aeruginosa* or was a result of the killing of MAB (such as eDNA). In line with this, DNase degradation experiments indicated that eDNA likely plays a role in our dual-species phenotype, although it did not completely abrogate the increase in biofilm formation.

Because DNase treatment was unable to completely prevent our dual-species phenotype, we also wanted to determine if the known exopolysaccharides of *P.* aeruginosa, alginate, Psl, or Pel, were involved. In this study we included both mucoid and non-mucoid strains of *P. aeruginosa*, both of which showed a similar increase in biofilm biomass. The increase in biomass for both mucoid and non-mucoid strains indicated that this was likely not an alginate dependent phenotype. PAO1 is known to produce primarily Psl but can also produce Pel. We found that disruption of Psl alone was not sufficient to prevent the increase in biofilm formation in the presence of MAB at the 24-hour time point. However, overall biofilm biomass was decreased compared to wild type in this strain regardless of condition. In contrast, disruption of Pel prevented the increase in biofilm with MAB present at the 24-hour time point, indicating that Pel is likely primarily responsible for this increase, particularly at earlier time points. Interestingly, the disruption of Pel also resulted in the emergence of an increase in the dual-species biofilm at the 72-hour time point, which is not seen in the parent strain. This might indicate that Pel is playing a large role in our biofilm phenotype at earlier time points, but Psl becomes more important as the biofilm matures. Disruption of both polysaccharides completely abrogated our dual-species phenotype. However, this strain showed very little biofilm formation regardless of MAB presence. Although mPA08-31 is a mucoid isolate, Congo red assays indicated that it also produces Pel, making it unsurprising that we saw our phenotype for this isolate. We also saw that PA14, which makes Pel, but is missing the Psl locus shows an increase in biofilm production with MAB present. FRD1, an alginate overproducer known to express smaller amounts of Psl and Pel showed the least amount of cooperativity, consistent with our proposed mechanism of a Pel-dependent increase in biofilm matrix with Psl participating likely in a redundant fashion.

Our results indicate that both Pel and eDNA are involved in the dual species increase in biofilm biomass, particularly at earlier time points. Direct ionic binding between eDNA and Pel has been demonstrated, and interactions between these molecules are thought to directly contribute to *P. aeruginosa* biofilm structure and rheology^66,69,70^. Although we did not directly investigate any changes in the structural interactions between these two molecules in the dual-species biofilm of MAB and *P. aeruginosa,* it is possible that the biofilm structure is being strengthened by interactions between the Pel derived from *P. aeruginosa* and the eDNA from MAB, providing a mechanism by which both species might show increased persistence in a host environment despite direct antagonism. This is also consistent with reports that these two organisms do not correlate with initial acquisition but do result in more severe and recalcitrant infections^52–54^.

Both Pel and eDNA, individually and when bound, are known to contribute to the of *P. aeruginosa* biofilms to aminoglycosides, including the CF relevant antibiotic tobramycin^66–68,71^. We therefore hypothesized that this is the mechanism by which co-culture with MAB is increasing tolerance to tobramycin in our dual-species biofilms. However, this could also be mediated by a variety of other mechanisms, including changes in metabolism or increased expression of efflux pumps. We did not see these systems increase in the RNA-sequencing analysis, although this was a later time point than what we used for the tobramycin assays, and this might have been picked up by studying transcriptomic changes at a variety of time points. As this is an important topic for patient outcomes, particularly for pwCF for which inhaled tobramycin is a part of standard treatment regimens, further investigation into the basis for this increase antibiotic tolerance would be warranted.

In our transcriptomic analysis we saw a nearly uniform decrease in the *amiEBCRS* operon of *P. aeruginosa* during co-culture with MAB. This operon encodes the aliphatic amidase AmiE along with its regulatory elements, which is responsible for the hydrolysis of short-chain aliphatic amides to their corresponding organic acids, a process conserved in all kingdoms of life^72^. Mycobacterial species have long been known to harbor a similar operon and produce functional aliphatic amidases^73,74^, therefore possibly negating the necessity of this system for *P. aeruginosa* during co-culture. The small non-coding RNA *amiL* encoded on this operon is negatively regulated by both *las* and *rhl* systems and is known to regulate a wide variety of virulence phenotypes. Deletion of AmiL increases elastase activity, pyocyanin production, and relative biofilm production^75^. AmiE overexpression in PA14 causes a decrease in biofilm thickness and twitching motility and a likewise decrease in molecules involved in virulence including HCN and pyocyanin^76^. Lastly, AmiC is responsible for biofilm dispersal after exposure to human C-type natriuretic peptide, again indicating an inverse relationship to biofilm formation in *P. aeruginosa*^77^. Ultimately, these studies indicate that the amidase operon is a negative regulator of many virulence phenotypes, and its downregulation is likely responsible for the increase in biofilm formation seen here and may be associated with a variety of other clinically relevant virulence phenotypes.

Although this study included several implications for further research and treatment options, it was limited by its restriction to an *in vitro* biofilm model in a rich medium. Further studies in more relevant conditions for CF disease, including a cell culture model or culture in synthetic cystic fibrosis sputum medium (such as SCFM2) are warranted to confirm that similar interactions are occurring in the context of CF.

Ultimately, we have demonstrated that co-culture of two relevant late-stage CF pathogens, *P. aeruginosa* and MAB, contributes to disease-relevant phenotypes, including an increase in biofilm formation and antimicrobial tolerance. We have also demonstrated that eDNA and Pel, two components of the biofilm matrix known to contribute to infection recalcitrance are involved in our increase in biofilm formation. Lastly, we proposed a novel mechanism by which these two organisms may interact, with MAB inducing a downregulation of the aliphatic amidase of *P. aeruginosa,* possibly leading to the downstream changes in biofilm formation.

## METHODS

### Strains and growth conditions

*P. aeruginosa* strains used in this study include PAO1 (provided by D. Wozniak, Ohio State University) and PA14, two widely used non-mucoid model strains, mPA08-31, a mucoid clinical isolate (provided by S. Birket, University of Alabama at Birmingham), and FRD1, a model mucoid strain. *P. aeruginosa* mPA08-31^GFP+^ and PAO1^GFP+^ were constructed by transforming parent strains with plasmid pSMC2 (provided by G. O’Toole, Dartmouth College). PAO1Δ*pelA*, PAO1Δ*pslA*, PAO1ΔΔ*pslApelA* were obtained from the Manoil Lab mutant library collection. *P. aeruginosa* strains were maintained on *Pseudomonas* isolation agar (PIA: 20g peptone, 1.4g magnesium chloride, 10g potassium sulfate, 25mg Irgasan, 21.1g agar, 12.5g LB per liter), were subsequently cultured in Lysogeny broth (LB: 10g tryptone, 5g yeast extract, 10g sodium chloride per liter), and were incubated at 37°C shaking at 250rpm.

*M. abscessus* strains used in this study include ATCC19977 rough and smooth isotypes, obtained from the American Type Culture Collection (ATCC). MAB(S)^mCherry+^ and MAB(R)^mCherry+^ were constructed by transforming parent strains with plasmid pMSP12::mCherry, obtained from Addgene (#30169), using previously described methods^78^. Each morphotype was transformed separately, and we saw no evidence of conversion in this strain (Supplementary Figure 1C). *M. abscessus* strains were maintained on 7H10 agar (Remel) with 10% oleic acid-albumin-dextrose-catalase (OADC) supplement (BD) and glycerol, were subsequently cultured in 7H9 medium (Remel) with 0.05% Tween, 10% albumin-dextrose-catalase (ADC) supplement (BD), and glycerol, and were incubated at 37°C shaking at 80rpm.

### Static biofilm formation assays

*In vitro* biofilm assays were performed according to an established microtiter assay protocol with some modifications^79^. Overnight cultures of *P. aeruginosa* were sub-cultured 1:100 in LB and grown up to mid-log phase (OD at 600nm = 0.4-0.6). Broth stocks of MAB grown to stationary phase in 7H9 (+Tween, OADC) were subcultured 1:100 in fresh media and grown overnight (37°C, 80 rpm) to midlog phase (OD at 600nm = 0.4-0.6). MAB was pelleted (5000g, 5 min) and resuspended in LB. Both species were inoculated either separately or together into a 96-well microtiter dish (Corning) in LB at a final density of ∼10^6^ CFU/mL for *P. aeruginosa* and ∼10^7^ for MAB to account for differences in growth rate (200μL/well). Biofilms were incubated without shaking at 37°C for the time specified for each assay.

To quantify the resulting biofilm, plates were washed, stained with 0.1% crystal violet, and dissolved in 30% acetic acid. Absorbance was first measured at 600nm prior to washing, and then 562 nm after crystal violet staining to quantify biofilm biomass using the Synergy HTX Multi-Mode Microplate Reader (BioTek). Biofilm formation is expressed as biomass normalized to growth (OD562/OD600) unless otherwise specified.

### Quantification of *P. aeruginosa* and MAB during co-culture

Biofilms were prepared as described above in 96-well microtiter dishes. To quantify colony forming units of each species, adherent biofilm cells were then washed two times with phosphate-buffered saline (PBS), scraped and resuspended in 200ul of PBS before serial dilution and selective plating using the track dilution method^80^ with a limit of detection at 10^2^. A selective plating scheme of MacConkey agar (Remel) to grow *P. aeruginosa* alone and Columbia CNA agar (BD) with 5% sheep’s blood (Hemostat Laboratories) to grow MAB was used as previously described^58,59^

### Confocal microscopy

Single- and dual-species biofilms were prepared as described above in a sterile eight-well treated μSlide (Ibidi) using fluorescently labeled strains *P. aeruginosa* PAO1^GFP+^, mPA08- 31^GFP+^, MAB(S)^mCherry+^ and MAB(R)^mCherry+^. Adherent biofilm cells were then washed two times with phosphate-buffered saline (PBS) and fixed using Fluoromount-G (Invitrogen). Confocal laser scanning microscopy (CLSM) was performed using a Nikon-A1R HD25 confocal laser microscope (Nikon) at 40X magnification. Images were acquired and processed using the NIS-elements 5.0 software. For analysis, images were first denoised using the Denoise.ai feature of NIS elements software. Image stacks were then imported into a biofilm quantification software, BiofilmQ,^81^ as ND2 files to preserve metadata. Image stacks were thresholded for each channel individually, differentiating between biofilm and background signal. To analyze properties of each biofilm volume with spatial resolution, BiofilmQ employs a cube-based method to define localization. Defined volumes from fluorescent images were dissected into individual pseudo-cell cubes, (side length is 3 vox, or 0.94um) for which parameter calculations were then performed. Metrics calculated included Mean Biofilm Volume (µm3/area), Mean Biofilm Height (µm/area), and Distance to Substrate (µm/voxel) for each 1024 x 1024 pixel (322µm x 322µm) substrate area, with substrate being the bottom surface of the 8-well slide. Three images were analyzed per biofilm, for at least 2 biological replicates.

### DNase treatment

For quantification of biofilm biomass with and without DNase treatment, single- and dual-species biofilms were grown as detailed above for 24, 48, or 72 hours. OD at 600nm was taken before supernatant was removed and replaced with DNase (25 U/mL) in PBS. Biofilms were incubated with DNase treatment at 37°C for 1 hour before performing a crystal violet assay as previously described. For confocal imaging, single- and dual-species biofilms were grown for 24 hours for PAO1^GFP+^ biofilms and 72 hours for mPA08-31^GFP+^ biofilms. Supernatant was removed and replaced with DNase (25 U/mL) in PBS. Adherent biofilm cells were then washed two times with phosphate-buffered saline (PBS) and fixed using Fluoromount-G (Invitrogen) before performing imaging and quantification as described above.

### Tobramycin susceptibility

For quantification of biofilm biomass with and without tobramycin treatment, single- and dual-species biofilms were grown as detailed above for 24 hours. Supernatant was then removed and replaced with tobramycin (50ug/mL) in PBS or a PBS control. Biofilms were incubated with tobramycin treatment at 37°C for 1 hour before performing a crystal violet assay as previously described. For confocal imaging, single- and dual-species biofilms were grown for 24 hours. Supernatant was removed and replaced with tobramycin (50ug/mL) in PBS. Adherent biofilm cells were then washed two times with phosphate-buffered saline (PBS) and fixed using Fluoromount-G (Invitrogen) before performing imaging and quantification as described above.

### RNA-sequencing and analysis

For single- and dual-species RNA-seq, biofilms were prepared as described above with some modifications. Overnight cultures of *P. aeruginosa* were sub-cultured 1:100 in LB and grown up to mid-log phase (OD at 600nm = 0.4-0.6). Broth stocks of MAB grown to stationary phase in 7H9 (+Tween, OADC) were subcultured 1:100 in fresh media and grown overnight (37°C, 80 rpm) to midlog phase (OD at 600nm = 0.4-0.6). MAB was pelleted (5000g, 5 min) and resuspended in LB. Both species were inoculated either separately or together into a 6-well microtiter dish (Corning) in LB at a final density of ∼10^6^ CFU/mL for *P. aeruginosa* and ∼10^7^ (3mL/well). Biofilms were incubated without shaking at 37°C for 48 hours before being washed twice with PBS. Adherent cells from 3 wells were pooled for each sample. Samples were pelleted and resuspended in TRIzol reagent (Invitrogen) before bead beating (0.1mm silica) to lyse cells. RNA was extracted according to a standard protocol^82^.

Isolated RNA was sent to SeqCenter (Pittsburgh, PA) for rRNA and tRNA depletion, sequencing, and analysis. Reads were aligned to either the PAO1 or mPA08-31 genome as appropriate using HISAT2 (v. 2.2.0). Read counts were obtained via Subread featureCounts (v.2.0.1), and differential expression analysis was performed with DESeq2 with an adjusted P-value cutoff of <0.05.

### RT qPCR

Overnight cultures of *P. aeruginosa* were sub-cultured 1:100 in LB and grown up to mid-log phase (OD at 600nm = 0.4-0.6). Broth stocks of MAB grown to stationary phase in 7H9 (+Tween, OADC) were subcultured 1:100 in fresh media and grown overnight (37°C, 80 rpm) to midlog phase (OD at 600nm = 0.4-0.6). MAB was pelleted (5000g, 5 min) and resuspended in LB. Both species were inoculated either separately or together into a 6-well microtiter dish (Corning) in LB at a final density of ∼10^6^ CFU/mL for *P. aeruginosa* and ∼10^7^ (3mL/well). Biofilms were incubated without shaking at 37°C for 48 hours before being washed twice with PBS. Adherent cells from 3 wells were pooled for each sample. Samples were pelleted and resuspended in TRIzol reagent (Invitrogen) before RNA extraction using the Direct-zol DNA/RNA Miniprep kit (Zymo).

Extracted RNA was converted to cDNA via the iScript cDNA Synthesis Kit (Bio-rad) and RT-qPCR was performed using the CFX Opus 96 Real-Time PCR System, and probes specific for the genes of interest and with no significant homology to the MAB genome. Gene expression was normalized to 16S expression via species-specific probes. iTaq Universal SYBR Green Supermix was used according to the manufacturer’s directions for cycling conditions (Biorad). All samples were run in duplicate. Transcript measures were normalized relative to 16S levels from the same sample. Relative quantification of gene expression was determined using the comparative cycle threshold (CT) method (2^-ΔΔCT^).

### Statistical analyses

Unless otherwise noted, graphs represent sample means ± standard error of the mean (SEM). For nonparametric analyses, differences between groups were analyzed by Kruskal-Wallis test with the uncorrected Dunn’s test for multiple comparisons. For normally distributed data sets (as determined by Shapiro-Wilk normality test) a one-way analysis of variance (ANOVA) was used with Tukey’s multiple-comparison test. Those experiments with two factors (for example, CFU changes over time) were analyzed via two-way ANOVA. Outliers were detected via the ROUT method (Q, 1%) and excluded from the analysis. For all relevant statistical analyses, an alpha value of 0.5 was used and significance was denoted as follows: *p <0.05, **p < 0.01, ***p < 0.001, ****p < 0.0001. For viable colony counts, data was transformed (Y=log_10_(Y)) to achieve normally distributed data appropriate for use with a Two-way ANOVA. All statistical tests were performed using GraphPad Prism 10 (San Diego, CA).

## DATA AVAILABILITY

The raw sequencing data from the current study are available in the sequence read archive (SRA) under the bioproject PRJNA1073745.

## ACKNOWLEDGEMENTS

This study was funded by the Cystic Fibrosis Foundation (MCDANI23F0) awarded to M.M. and an NIH R35 (GM142748) awarded to J.S. SE was previously funded by the NIDCR/Dental Academic Research Training Program (T90DE022736) and is now supported by a NIH/NIDCR National Research Service Award (1F31DE034608). E.P. was supported by the UAB Cystic Fibrosis Summer Research Program. J.B. was supported by a NIH/NIDCR National Research Service Award (F31DE031508). The funders played no role in study design, data collection, analysis and interpretation of data, or the writing of this manuscript.

We thank Shawn Williams and Robert Grabski at the University of Alabama at the Birmingham High Resolution Imaging Facility for their assistance with the Nikon A1 confocal microscope and imaging analysis. Daniel Wozniak (Ohio State University) and Susan Birket (University of Alabama at Birmingham) provided bacterial strains for this study.

## AUTHOR CONTRIBUTIONS

M.M. and J.S. conceived and planned the experiments, with J.S. supervising the work. M.M., E.P., and S.E., carried out experiments. M.M., E.P., S.E., J.B., and J.S. contributed to the interpretation of the results. M.M. took the lead in designing figures and writing the manuscript, with significant contributions from J.S. during editing. All authors provided critical feedback and helped shape the research, analysis and manuscript.

