## Supplemental File 1 for "*Mycobacterium abscessus* promotes *Pseudomonas aeruginosa* biofilm formation and antibiotic tolerance"

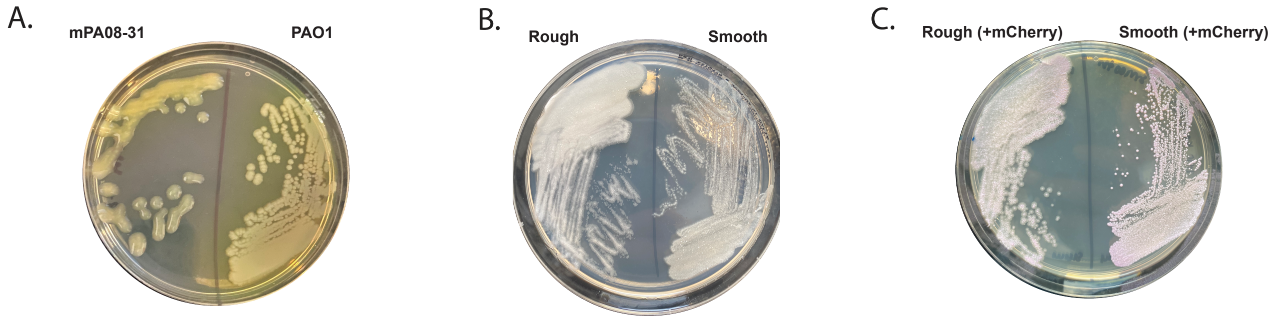


**Supplementary Figure 1. Morphotypes of strains used**
A) Image of non-mucoid *P. aeruginosa* PAO1 and mucoid *P. aeruginosa* mPA08-31 plated on LB agar. B) Image of rough and smooth morphotypes of *M. abscessus* ATCC 19977 plated on 7H10 agar. C) Image of rough and smooth morphotypes of *M. abscessus* ATCC 19977 (mCherry+) plated on 7H10 agar (+kanamycin 50μg/mL).

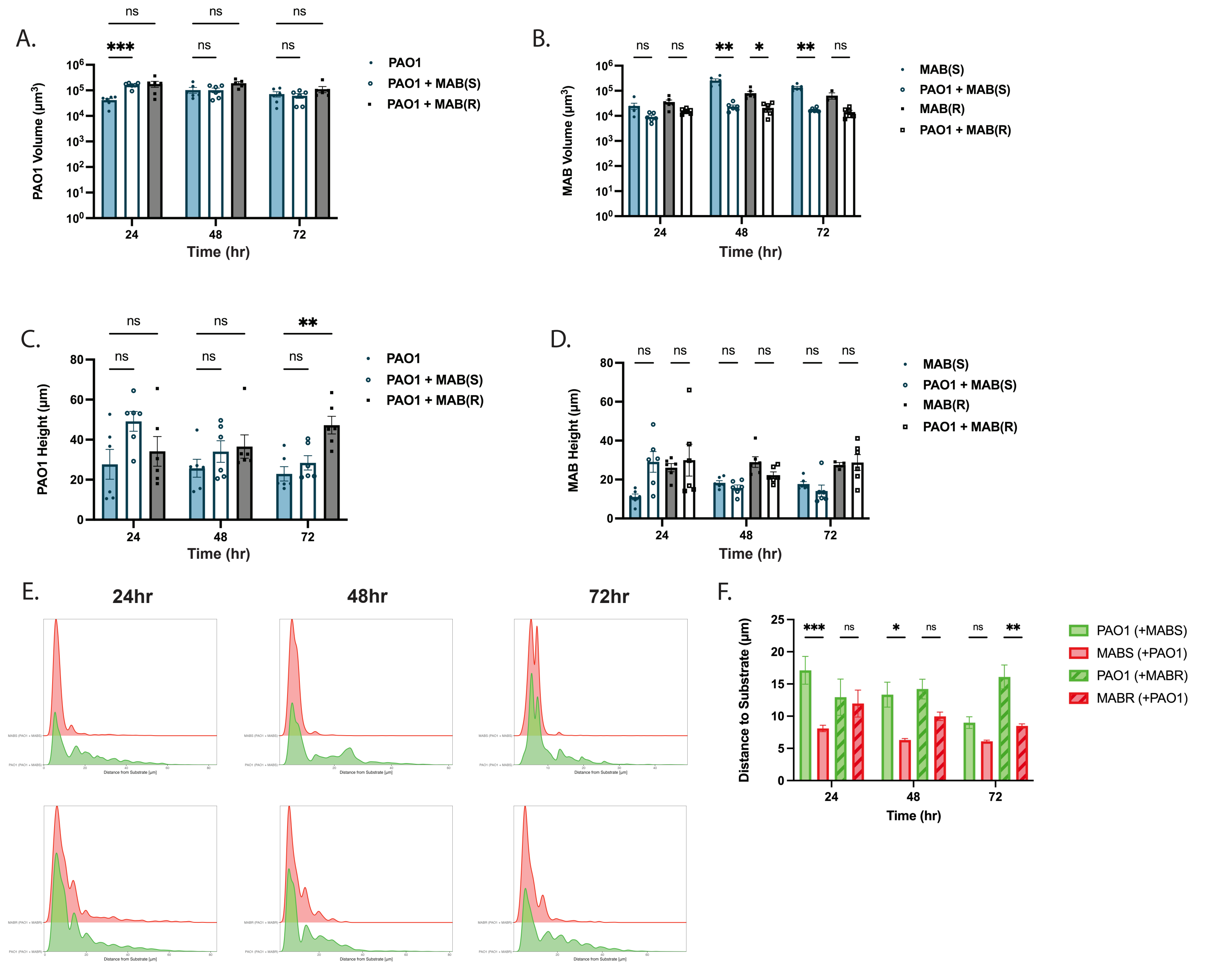


**Supplementary Figure 2. Imaging of dual species *P. aeruginosa* PAO1 and MAB biofilms**
Fluorescently labeled *M. abscessus* ATCC19977 smooth and rough morphotypes (mCherry+) were co-cultured with *P. aeruginosa* PAO1 (GFP+) in LB medium in an 8-well µSlide for 24, 48, or 72 hours at 37 °C (n = 2 biological replicates, 3 technical). A) Volume and B) height of *P. aeruginosa* PAO1 and C) volume and D) height of *M. abscessus* in single- and dual-species biofilms were quantified via BiofilmQ. One-way ANOVA with Tukey's multiple comparisons test. E) Histograms of pixel distribution of *P. aeruginosa* PAO1 and *M. abscessus* distance to substrate at each timepoint for rough and smooth morphotypes. F) Mean distance to substrate summarized. (n = 2 biological replicates, 3 technical).

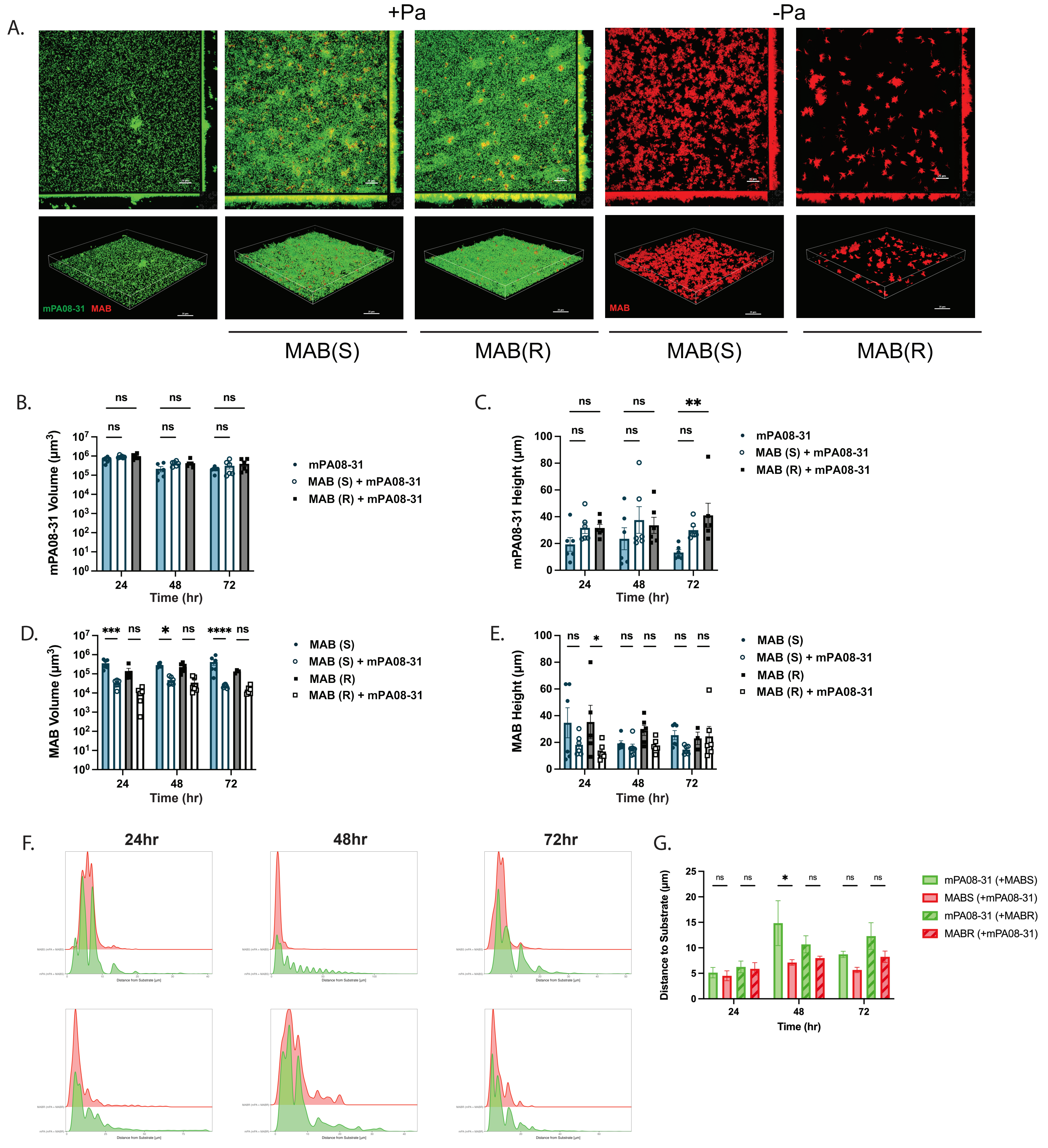


**Supplementary Figure 3. Imaging of dual species *P. aeruginosa* mPA08-31 and MAB biofilms**
Fluorescently labeled *M. abscessus* ATCC19977 smooth and rough morphotypes (mCherry+) were co-cultured with *P. aeruginosa* mPA08-31 (GFP+) in LB medium in an 8-well µSlide for 24 hours at 37 °C (n = 2 biological replicates, 3 technical). A) Structural composition of single- and dual-species biofilms was evaluated via confocal imaging at 40X magnification. B) Volume and C) height of *P. aeruginosa* mPA08-31 and D) volume and E) height of *M. abscessus* in single- and dual-species biofilms were quantified via BiofilmQ. One-way ANOVA with Tukey's multiple comparisons test. F) Histograms of pixel distribution of *P. aeruginosa* PAO1 and *M. abscessus* distance to substrate at each timepoint for rough and smooth morphotypes. G) Mean distance to substrate summarized. (n = 2 biological replicates, 3 technical).

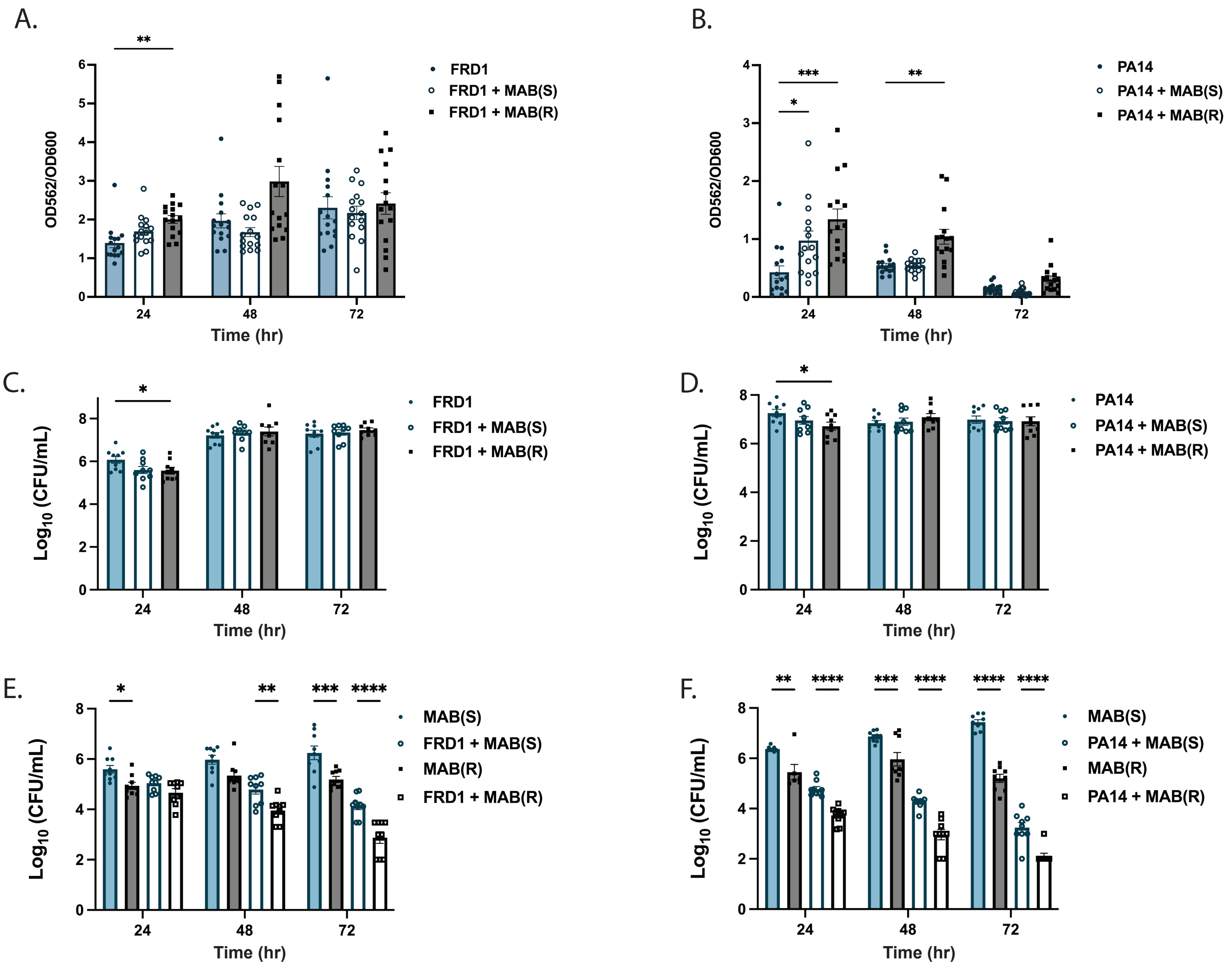


**Supplementary Figure 4. Dual species biofilms with *P. aeruginosa* strains PA14 and FRD1**
*M. abscessus* ATCC 19977 smooth and rough morphotypes were co-cultured with either of two *P. aeruginosa* strains (FRD1 or PA14) in LB medium in a 96-well plate for 24, 48, or 72 hours at 37 °C. Biofilm biomass was then measured using crystal violet staining for A) FRD1 or B) PA14 during single- and co-culture (n = 3 biological replicates, 6 technical). Two-way ANOVA with Tukey's multiple comparisons test. Viable colony forming units of *P. aeruginosa* C) FRD1 or D) PA14 during single- and co-culture, and of M. abscessus during single- and co-culture with E) FRD1 and F) PA14 were measured (n = 3 biological replicates, 3 technical). Two-way ANOVA with Tukey's multiple comparisons test.

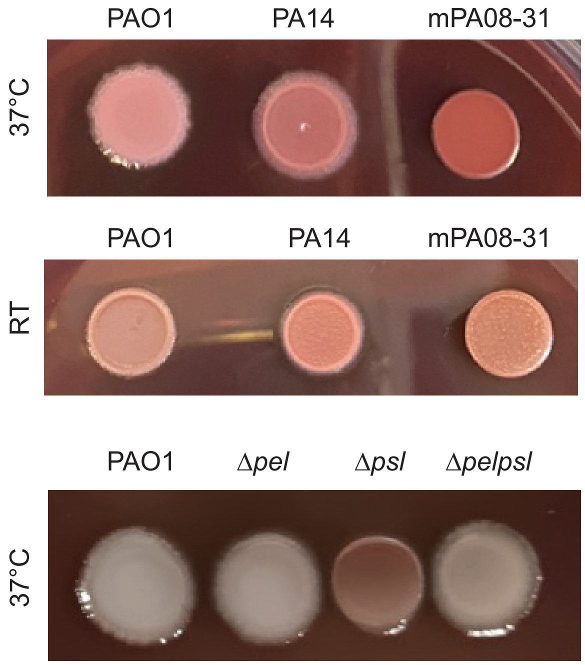


**Supplementary Figure 5.** ***P. aeruginosa*** **morphology on Congo Red plates**
*P. aeruginosa* strains PAO1, PA14, and mPA08-31 were grown on Congo red plates (LB, no NaCl, 40μ g/mL Congo red and 20μg/mL Coomassie blue) at either 37 °C or room temperature for 48 hours. *P. aeruginosa strains* PAO1, PAO1Δ*pel*, PAO1Δ*psl*, and PAO1Δ*pelpsl* were grown on Congo red plates at 37 °C for 48 hours.

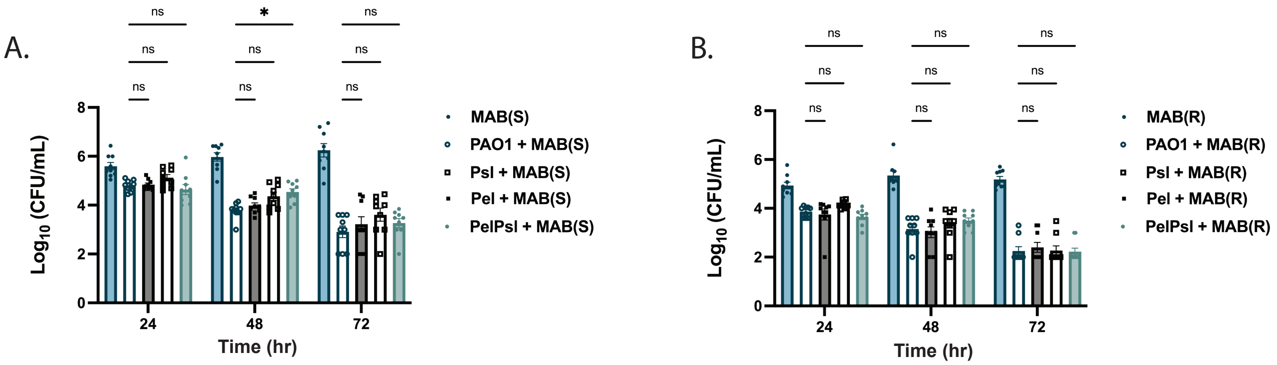


**Supplementary Figure 6. *P. aeruginosa* mutants show no killing defect against MAB**
*M. abscessus* ATCC 19977 smooth and rough morphotypes were co-cultured with *P. aeruginosa* strains PAO1, PAO1Δpel, PAO1Δpsl, and PAO1Δpelpsl in LB medium in a 96-well plate for 24, 48, or 72 hours at 37 °C. Viable colony forming units of *M. abscessus* A) smooth and B) rough morphotypes in single- and dual-species biofilms were measured. (n = 3 biological replicates, 3 technical). Two-way ANOVA with Tukey's multiple comparisons test.

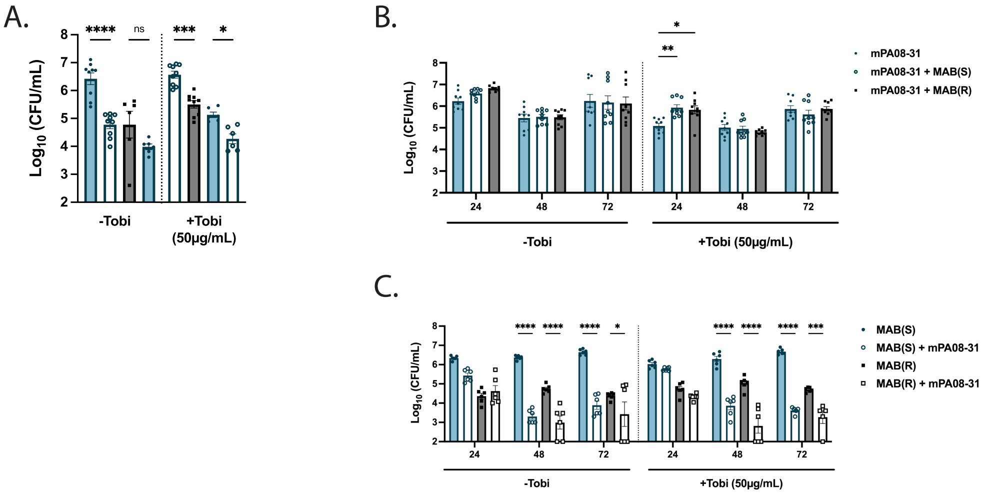


**Supplementary Figure 7. Tobramycin treatment does not affect killing of MAB by *P. aeruginosa***
*M. abscessus* ATCC 19977 smooth and rough morphotypes were co-cultured with PAO1 or in LB medium in a 96-well plate for 24, 48, or 72 hours at 37 °C. Biofilms were then treated for 1 hour with tobramycin (50 µg/mL) or 1X PBS. A) Viable colony forming units of *M. abscessus* in single- or dual-species biofilms with *P. aeruginosa* PAO1 were measured. Viable colony counts of B) *P. aeruginosa* mPA08-31 and C) *M. abscessus* during single- and dual-species culture with and without tobramycin treatment were measured via differential plating at all time points. (n = 3 biological replicates, 3 technical). One-way ANOVA with Tukey's multiple comparisons test.

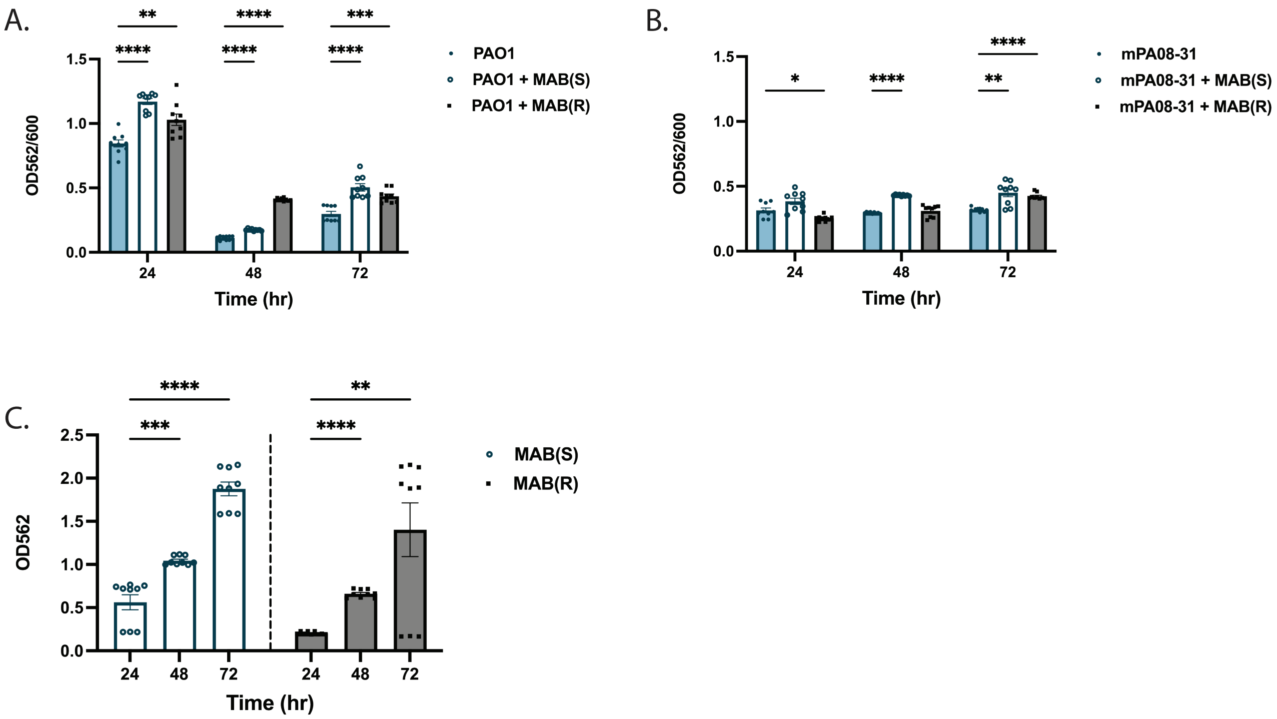


**Supplementary Figure 8. Adaptation of experimental setup to a 6-well plate format for RNA-sequencing**
*M. abscessus* ATCC 19977 smooth and rough morphotypes were co-cultured with either of two *P. aeruginosa* strains (PAO1 or mPA08-31) in LB medium in a 6-well plate for 24, 48, or 72 hours at 37 °C. Biofilm biomass was then measured using crystal violet staining for A) PAO1 or B) mPA08-31 during single- and co-culture. C) Development of single-species biofilms of *M. abscessus* over time as measured via crystal violet assay (OD562). Normalization was not used due to relatively low OD600 readings in comparison to P. aeruginosa. (n = 3 biological replicates, 3 technical). Two-way ANOVA with Tukey's multiple comparisons test.
cal). One-way ANOVA with Tukey's multiple comparisons test.

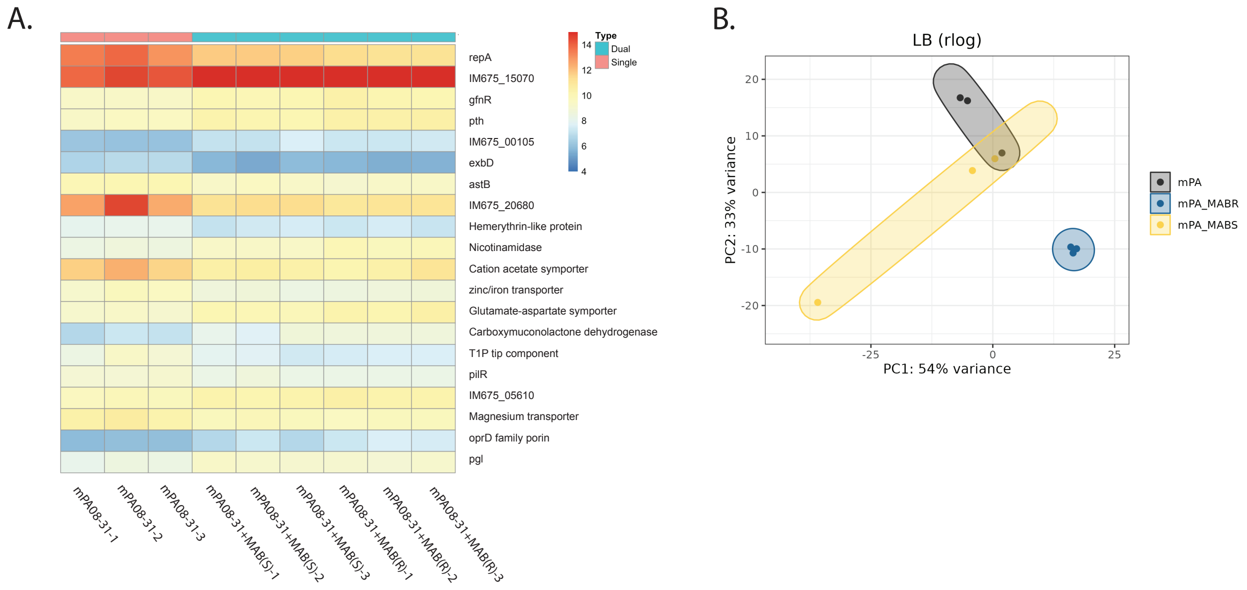


**Supplementary Figure 9. RNA sequencing of *P. aeruginosa* mPA08-31 biofilms with and without MAB**

*M. abscessus* ATCC 19977 smooth and rough morphotypes were co-cultured with *P. aeruginosa* mPA08-31 in LB medium in a 6-well plate for 48 hours at 37 °C. Adherent cells were pooled (3 per sample) for RNA extraction and sequencing (n = 3). A) Heatmap of differentially expressed genes during mPA08-31 single or dual-species biofilms determined via differential expression analysis using DESeq2. B) Principal-component analysis of samples based on RNA sequencing data, colored by infection group.

**Supplementary Data 1. Gene counts mapped to *M. abscessus* ATCC19977 in RNA-sequencing experiments**

**Supplementary Data 2. Gene counts mapped to *P. aeruginosa* PAO1 in RNA-sequencing experiments**

**Supplementary Data 3. Gene counts mapped to *P. aeruginosa* mPA08-31 in RNA-sequencing experiments**

**Supplementary Data 4. Differential expression of genes from *P. aeruginosa* PAO1 in single- and dual-species biofilms**

**Supplementary Data 5. Differential expression of genes from *P. aeruginosa* mPA08-31 in single- and dual-species biofilms**
